# HapNet: a new Python package for automated population-aware haplotype network analysis and visualization

**DOI:** 10.64898/2026.02.18.706154

**Authors:** Andrew A. Davinack

**Affiliations:** Department of Biological, Chemical, and Environmental Sciences, Wheaton College, Massachusetts, Norton, MA 02766

## Abstract

Haplotype networks are widely used in population genetics and phylogeography to visualize genealogical relationships among DNA sequences and to infer population structure, historical connectivity, and demographic processes. Existing software for haplotype network construction relies primarily on interactive graphical interfaces, which limits reproducibility, automation, and integration into modern bioinformatic workflows. Here, I introduce HapNet, an open-source Python package that enables automated construction, visualization, and summarization of haplotype networks directly from aligned FASTA files. HapNet is the first Python-native package designed specifically for automated, population-aware haplotype network construction and visualization from aligned FASTA files. HapNet implements a minimum-spanning-tree approach based on Hamming distances among haplotypes and incorporates population metadata encoded in sequence headers to produce population-aware network visualizations in which shared haplotypes are represented as pie charts and node sizes scale with haplotype frequency. In addition to a publication-ready network, HapNet generates machine-readable tabular output describing haplotype composition, population membership, and shared versus private haplotypes, facilitating downstream statistical analysis and reproducibility. Here, HapNet’s utility is demonstrated using mitochondrial DNA sequences from the shell-boring polychaete worm *Polydora neocaeca*, illustrating how the software reveals patterns of population connectivity and haplotype sharing. HapNet provides a reproducible, scriptable alternative to existing graphical tools and is freely available via the Python Package Index and GitHub.

## Introduction

Haplotype networks provide an intuitive graphical representation of genealogical relationships among DNA sequences and are widely used in population genetics, phylogeography, and molecular ecology to infer population structure, historical connectivity, and demographic change (Posada and Crandall 2001; Mardulyn 2012). By representing mutational steps among distinct haplotypes, such networks offer an alternative to strictly bifurcating phylogenetic trees when analyzing intraspecific genetic variation. Programs such as TCS and PopART have become standard tools for constructing haplotype networks, particularly for mitochondrial and chloroplast DNA data (Clement et al. 2000; Leigh and Bryant 2015). However, these programs are primarily graphical and interactive, making it difficult to automate analyses, ensure reproducibility, or integrate haplotype network inference into larger bioinformatic pipelines.

As genetic datasets grow in size and complexity, and as reproducibility becomes an increasingly important component of population genetic research, there is a need for haplotype network software that can be executed from the command line, integrated into scripted workflows, and produce machine-readable outputs alongside graphical visualization. Furthermore, many population genetic studies require explicit representation of population membership, yet existing tools often require manual assignment of individuals into populations within graphical interfaces, introducing opportunities for error and limiting scalability.

To address these limitations, HapNet was developed – a lightweight Python-based software package for automated population-aware haplotype network analysis. HapNet reads aligned FASTA files in which population identity is encoded in the sequence headers, constructs color coded haplotype networks using a minimum-spanning-tree framework, and produces both publication-quality networks and tabular outputs summarizing haplotype and population structure. Although there are R packages such as *pegas* which provide functions for haplotype network inference, they require custom scripting to produce population-aware network visualizations. HapNet is the first Python package to provide a fully-automated, end-user workflow for haplotype network visualization and summarization.

### Software design and implementation

HapNet is implemented in Python and distributed as an installable package via the Python Package Index (PyPI). The software operates from the command line and accepts a single aligned FASTA file as input. Population identity is inferred from the final underscore-delimited token in each sequence header, allowing users to encode population information directly within standard FASTA files without the need for auxiliary metadata files (Fig. 1).

**Figure 1.**
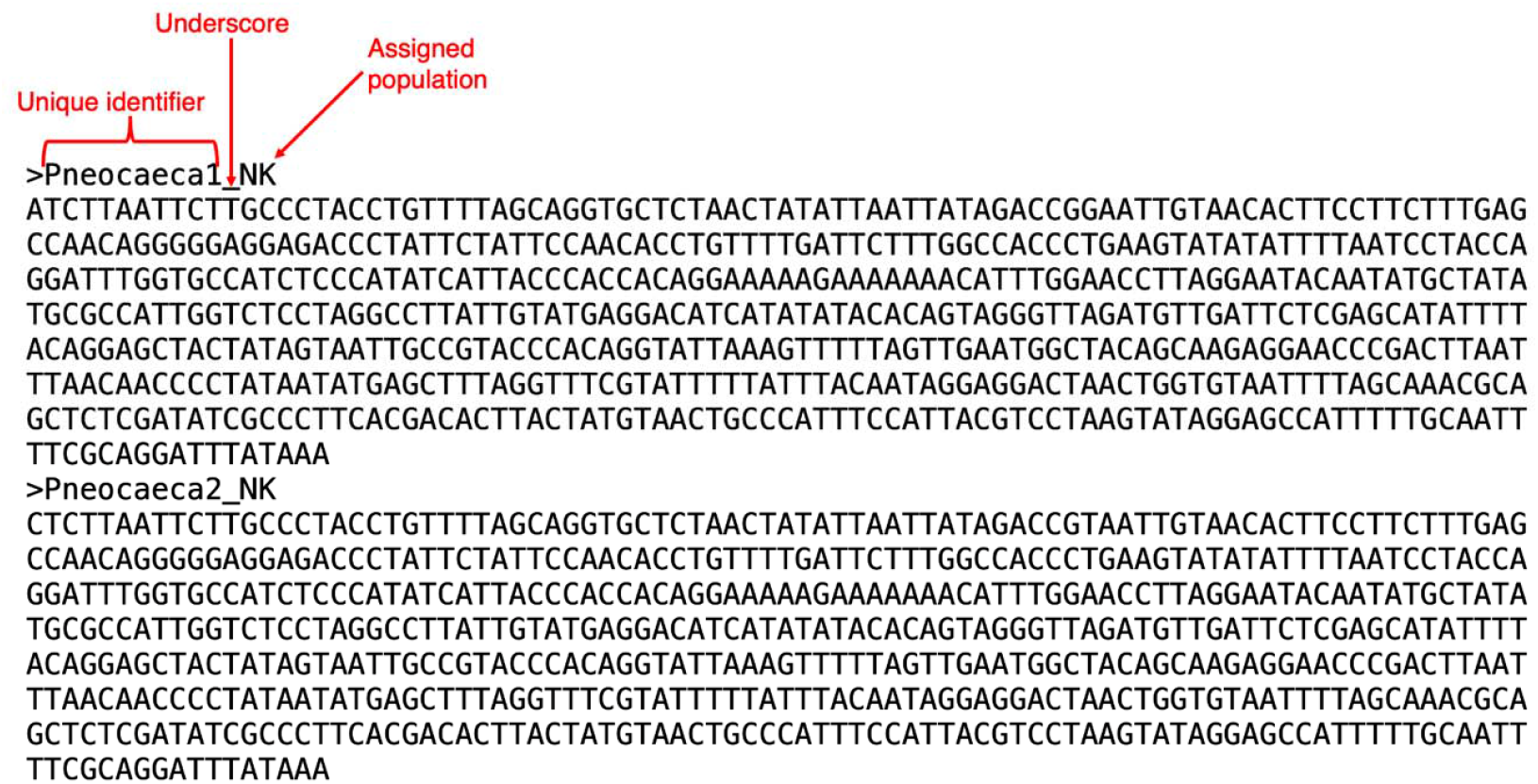
Example of HapNet input format for *Polydora neocaeca*. Population identity is encoded directly in FASTA sequence headers using an underscore-delimited suffix. Each header consists of a unique individual identifier followed by an underscore and a population code (e.g., Pneocaeca1_NK which refers to a single sample from Nantucket Island population).

The HapNet workflow proceeds in four main steps. First, sequences are read from the FASTA file and grouped by identical nucleotide strings to define haplotypes. Each haplotype stores a list of contributing individuals and their associated populations. Second, pairwise Hamming distances among haplotypes are computed across the aligned sequences, and a complete weighted graph is constructed. Third, a minimum-spanning tree (MST) is inferred from this graph, producing a parsimonious network connecting all haplotypes. Finally, the network is laid out and visualized using a force-directed algorithm, with node sizes called to haplotype frequency and node colors or pie-chart sectors indicating population composition. Mutational distances between haplotypes are displayed as tick marks on network edges.

In addition to the graphical output, HapNet writes a set of tab-delimited log files describing haplotype sequences, individual membership, population composition, and the identity of haplotypes shared among populations. These files provide a transparent and reproducible record of the network inference and enable downstream statistical analysis without manual transcription.

### Input and outputs

The primary input to HapNet is an aligned FASTA file in which all sequences have equal length. Population identity is encoded in the FASTA header as shown in Fig 1. The main output of HapNet is a network figure in PNG, PDF, or SVG format showing the haplotype network. Nodes are drawn with areas proportional to the number of individuals sharing that haplotype, and shared haplotypes are presented as pie charts with sectors colored by populations (colors are set by default). In addition, HapNet produces several .tsv files listing (i) all haplotypes and their frequencies, (ii) the assignment of individual sequences to haplotypes, (iii) haplotypes shared among populations, and (iv) summary statistics including the total number of haplotypes and populations. One explicit note about shared versus private haplotypes is that HapNet classifies haplotypes as either shared or private based on their population distribution. A shared haplotype is defined as one occurring in two or more populations, whereas a private haplotype is restricted to a single population, regardless of how many individuals carry it.

### Empirical example: *Polydora neocaeca* haplotypes

To illustrate the utility of HapNet, the software was applied to mitochondrial cytochrome c oxidase I (COI) sequences from the shell boring polychaete worm *Polydora neocaeca*. The species is a known parasite of bivalves, and was recently found inhabiting bay scallops on Nantucket Island (Davinack and Hill 2022). Because of their economic impact on shellfish aquaculture, *Polydora* worms have long been the focus of studies on dispersal and invasion. A dataset consisting of 18 samples in total were used for this example – these consisted of recently sequenced individuals from Nantucket Island (Massachusetts, USA) (N = 8), and previously published sequences from New York, USA (N = 1), Rhode Island, USA (N = 3) and South Africa (N = 6). For the latter three sites, the sequences are taken from a study by Malan et al. (2020).

Using an aligned FASTA file in which population identity was encoded in the sequence headers, HapNet automatically identified eight haplotypes of which one (H1) was shared between Rhode Island and Nantucket populations, whereas all other haplotypes were private to single populations. Haplotypes H2 and H8 were exclusive to South African samples and were separated from the main North America network by multiple mutational steps, indicating substantial COI divergence relative to haplotypes found in North America (Fig. 2). Notably, haplotype H2 was represented by multiple individuals from South Africa but was classified by HapNet as “private” because it was absent from all other populations, illustrating that HapNet distinguishes population sharing from within-population dependency.

**Figure 2.**
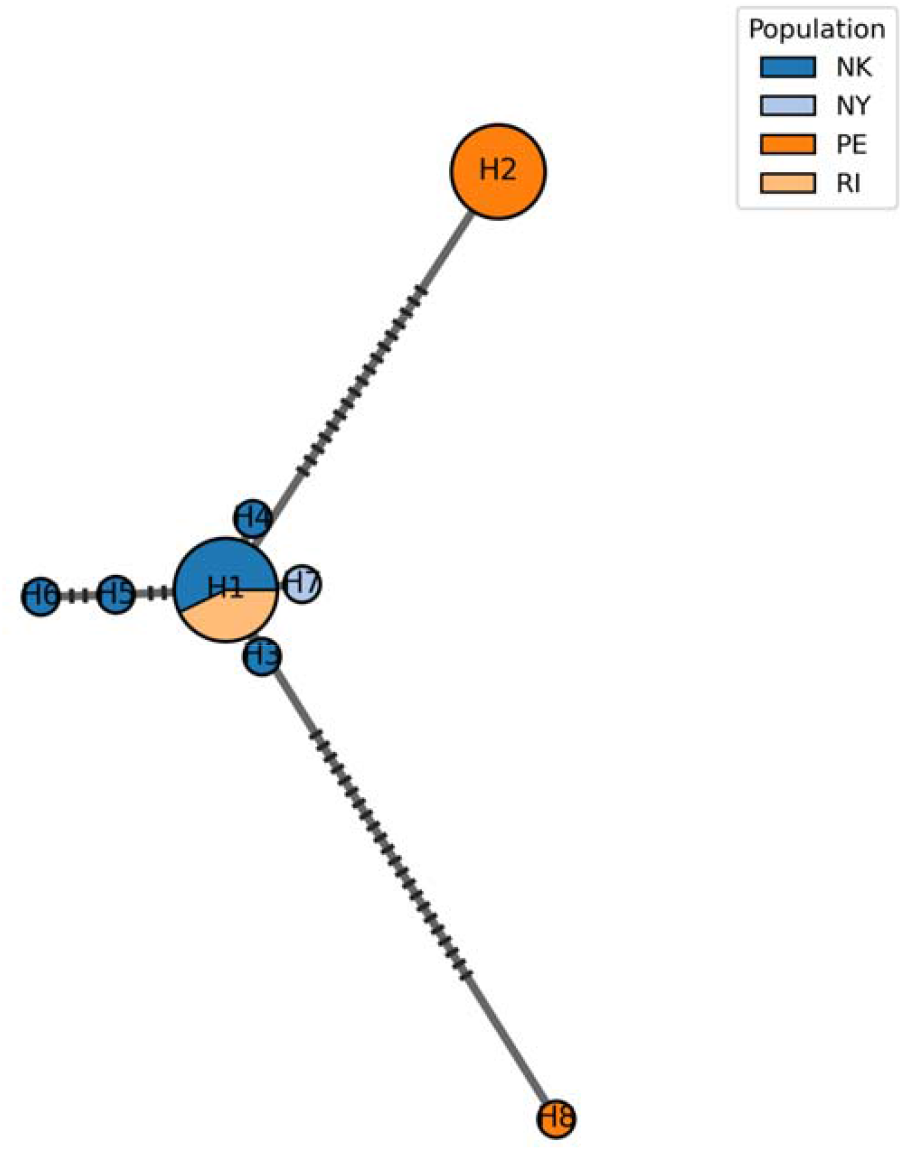
Population-aware haplotype network produced by HapNet. Circles represent unique haplotypes (H1 – H8) with circle area proportional to the number of sequences assigned to each haplotype. Colors within each circle indicate the proportional contribution of individuals from different populations, shown as pie charts when haplotypes are shared across populations. Solid edges connect haplotypes in a minimum-spanning tree based on Hamming distances among aligned sequences. Short black hash marks on edges indicate the number of mutational steps separating connected haplotypes. Population abbreviations correspond to geographic sampling locations (NK – Nantucket Island, NY – New York, USA, RI – Rhode Island, USA, PE – Port Elizabeth, South Africa).

Together, tables 1 – 4 provide a complete quantitative summary of haplotype composition, population membership, and cross-population sharing, documenting both the frequency and geographic distribution of all haplotypes identified in the dataset.

**Table 1.** Overall haplotype summary (from run1_summary.tsv)

| Metric | Value |
| --- | --- |
| Number of sequences | 18 |
| Number of haplotypes | 8 |
| Number of shared haplotypes | 1 |
| Number of private haplotypes | 7 |
| Maximum haplotype size | 7 |

**Table 2.**
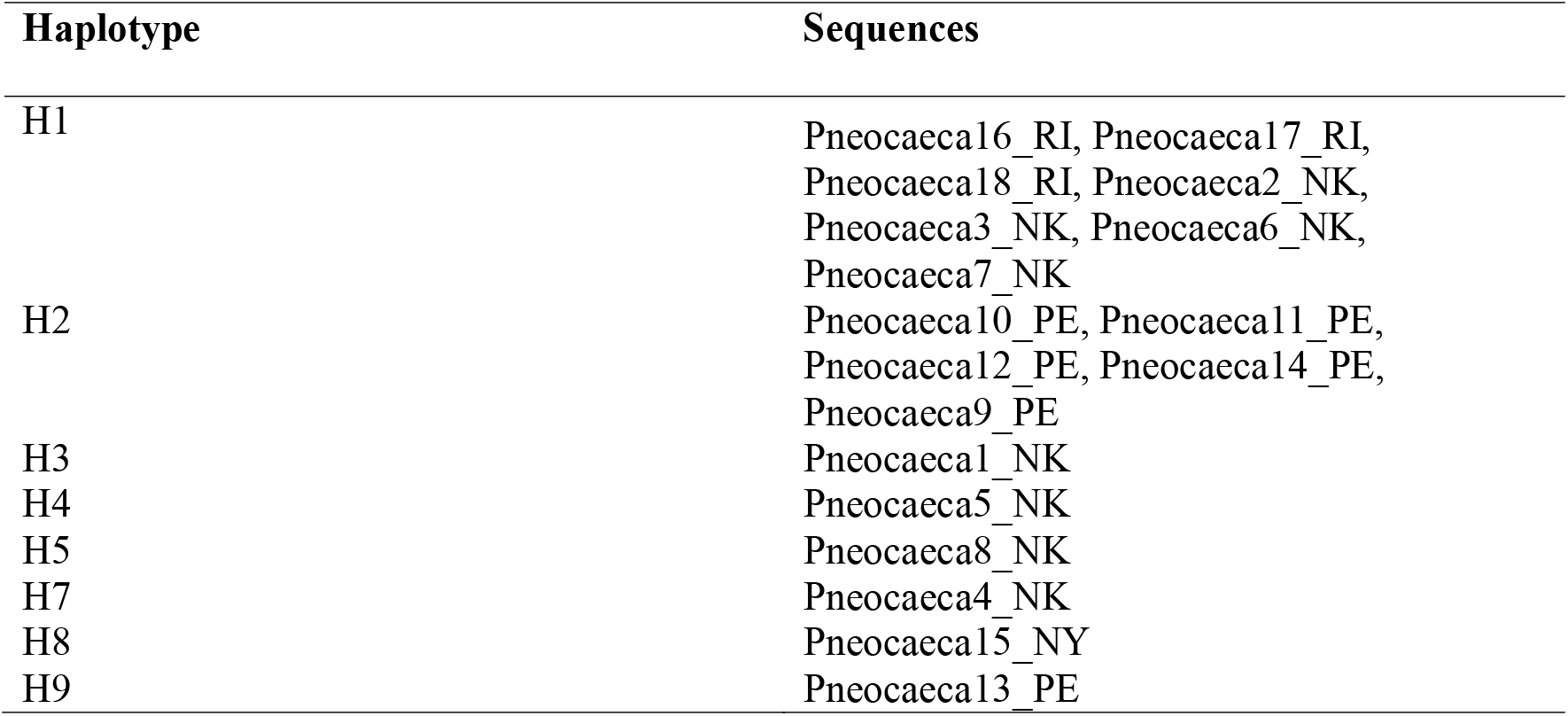
Haplotype membership (from run1_membership.tsv)

**Table 3.** Shared haplotypes across populations (from run1_shared_haplotypes.tsv)

| Haplotype | Populations | Counts by population |
| --- | --- | --- |
| H1 | NK, RI | NK: 4, RI: 3 |

**Table 4.** Haplotype composition and population distribution (from run1_haplotypes.tsv)

| hap_id | n_total | populations | counts_by_pop | sequences* |
| --- | --- | --- | --- | --- |
| H1 | 7 | NK, RI | NK:4, RI:3 | (not shown) |
| H2 | 5 | PE | PE:5 | (not shown) |
| H3 | 1 | NK | NK:1 | (not shown) |
| H4 | 1 | NK | NK:1 | (not shown) |
| H5 | 1 | NK | NK:1 | (not shown) |
| H6 | 1 | NK | NK:1 | (not shown) |
| H7 | 1 | NY | NY:1 | (not shown) |
| H8 | 1 | PE | PE:1 | (not shown) |

\*nucleotide sequences omitted for brevity.

## Availability and requirements

HapNet is freely available as open-source software. The package can be installed from the Python Package Index (https://pypi.org/project/hapnet/) using `pip install hapnet` and is hosted at https://github.com/parasiteguy/hapnet. A haplotype network can be generated from an aligned FASTA file by running `hapnet input.fasta –-out network.png –log-prefix run1`, which produces a network figure and accompanying tab delimited log files describing haplotype and population composition. An example dataset, documentation and source code are provided to allow users to test and extend the software. HapNet runs on all major operating systems (Windows, macOS, and Linux) using the command line, and requires Python version 3.9 or higher along with standard scientific Python libraries including Biopython, NumPy, SciPy, NetworkX and Matplotlib, which are all installed automatically when the package is installed.

